# Foraging theory and the propensity to be obese: an alternative to thrift

**DOI:** 10.1101/278077

**Authors:** Ulfat Baig, Lavanya Lokhande, Poortata Lalwani, Suraj Chawla, Milind Watve

## Abstract

The evolutionary origin of obesity is classically believed to be genetic or developmentally induced thrift, as an adaptation to ancestral feast and famine conditions. However, recently the thrift family of hypotheses have attracted serious criticism necessitating alternative thinking. Optimization of foraging behaviour is an important aspect of behavioural evolution. For a species evolved for optimizing nutritional benefits against predation or other foraging risks, reduction in foraging risk below a threshold dramatically increases the steady-state body weight. In modern life where feeding is detached from foraging, the behavioural regulation mechanisms are likely to fail resulting into escalation of adiposity. At a proximate level the signalling pathways for foraging optimization involve fear induced signal molecules in the brain including Cocaine and Amphetamine Regulated Transcript (CART) interacting with adiposity signals such as leptin. While leptin promotes the expression of the fear peptides, the fear peptides promote anorectic action of leptin. This interaction promotes foraging drive and risk tolerance when the stored energy is low and suppresses hunger and foraging drive when the perceived risk is high. The ecological model of foraging optimization and the molecular model of interaction of these peptides converge in the outcome that the steady state adiposity is an inverse square root function of foraging risk. The foraging optimization model is independent of thrift or insurance hypotheses, but not mutually exclusive. We review existing evidence and suggest testable predictions of the model. Understanding obesity simultaneously at proximate and ultimate levels is likely to suggest effective means to curb the obesity epidemic.

## Background

Classically the tendency to become obese which is increasingly observed globally in the human population is attributed to “thrift” and different versions of the thrifty gene and thrifty phenotype hypotheses abound in obesity literature (Watve & Diwekar-Joshi 2016). The central concept of thrift is the tendency to store fat during periods of abundance to be used as reserve to combat periods of starvation. The common explanation of the prevalent obesity epidemic by the thrift family of hypotheses is that the human system, evolved for a feast and famine environment, faces uninterrupted feast in the modern lifestyle which leads to escalation of adiposity. Recently the thrift family of hypotheses are under serious criticism (Baig *et al.* 2011; Speakman 2006; Vaag *et al.* 2012; Watve 2013; Wells 2003). While the logical framework that storage of fat during the time of food abundance can be adaptive when faced with somewhat predictable periods of food crunch is sound and similar phenomenon are known in animals, whether it applies to the prevalent obesity epidemic in humans is being questioned. The main line of criticism is based on the following questions.

i. Whether stone-age was really marked by feast and famine of sufficient intensity for thrift selection. When exactly in human history the feast and famine selection operated? Are obese individuals really better at surviving famines? Speakman (2006) argued that such harsh famines were uncommon in the human ancestry and therefore there may not be sufficient selective pressure for the evolution of thrift. Further there is no evidence that obese individuals have a better chance of surviving famine.
ii. Whether thrifty gene explains polymorphism or the variance in propensity observed in the population (Watve & Diwekar-Joshi 2016). Selection for thrift is unlikely to lead to stable polymorphism in a population (Baig *et al.* 2011) and therefore alternative explanations are needed for the population variability in the propensity to be obese.
iii. Where are obesity genes? The genome wide association studies have identified a large number of loci associated with obesity but they together explain a very small fraction of population variance in obesity parameters (Kilpeläinen *et al.* 2011; McCarthy & Zeggini 2009; Mutch & Clément 2006; Rankinen *et al.* 2006; Scott *et al.* 2007; Sladek *et al.* 2007; Thorleifsson *et al.* 2009). Therefore genetics is unlikely to be the major contributor to the population variance in the propensity to be obese. This finding undermines thrifty gene hypothesis, but not developmental plasticity for thrift. However, some other lines of evidence questions developmental origins of thrift as under.
iv. Can life-time programming based on foetal conditions be adaptive? Mathematical models show that for long lived species, foetal programming based on birth time conditions is unlikely to evolve unless there is a significant positive correlation between birth conditions and lifelong conditions since a mismatch can be detrimental (Baig *et al.* 2011; Bateson *et al.* 2004). As there is no detectable positive correlation between birth year and lifetime rainfall or other climatic conditions, foetal programming is unlikely to have evolved in anticipation of drought or famine. There is no evidence so far that individuals with intrauterine growth restriction (IUGR) are better adapted to starvation. Data from baboons as well as humans are on the contrary (Hayward *et al.* 2013; Lea *et al.* 2015).
v. Does foetal programming account for majority of obese and diabetic people? Although there is strong statistical evidence that individuals facing developmental constraints in critical growth phases are more likely to become obese, insulin resistant and develop type 2 diabetes (T2D) in adult life, the majority of adult obese or type 2 diabetics are not born with low birth weight (Boyko 2000; McCance *et al.* 1994). Therefore even if we accept that there is foetal programming for thrift, at the best it accounts for a limited fraction of obesity or type 2 diabetes in a population.
vi. Does thrift explain the systems level changes in obesity and diabetes? Obesity and insulin resistance is associated with a large number of changes in the different body systems and their functions (Koshiyama *et al.* 2008; Kulkarni *et al.* 2017; Watve & Yajnik 2007). The thrift hypotheses focus on energy homeostasis alone and offer little explanation as to why these diverse changes are associated with it. The classical assumption is that obesity inevitably leads to all other pathology and no evolutionary theory is needed. Interestingly in different species the physiological changes associated with obesity are different (Arinell *et al.* 2012; Bamford *et al.* 2016; De Koster & Opsomer 2012) and therefore evolutionary explanations for the complex network of changes associated with obesity is certainly needed (Watve and Diwekar-Joshi 2016, Kulkarni *et al.* 2017).
vii. What is the mechanism of thrift? The original suggestion of Neel was that high insulin levels were causal to obesity (Neel 1962). However, subsequently the thinking was turned upside down. Obesity was thought to be causal to insulin resistance which secondarily increased insulin levels. If the original mechanism of thrift proposed by Neel is no more believed, there should have been an alternative mechanism of thrift proposed. But so far no clear alternative proximate mechanism of thrift has been proposed and demonstrated. The body has a wide variety of mechanisms that regulate energy intake (Cummings & Overduin 2007; Friedman 2000; Morton *et al.* 2006; Speakman 2014; Stanley *et al.* 2005). Adequate thinking and investigation efforts have not gone into addressing the question how thrifty tendency overcomes these mechanisms. Some studies find that fat oxidation is impaired in obesity and this is a likely main contributing mechanism of human obesity (Oakes *et al.* 2013; Rogge 2009; Thyfault *et al.* 2004; Zunquin *et al.* 2009; Zurlo *et al.* 1990). This poses a paradox for the thrift family of hypotheses. Fat storage will be useful if it can be efficiently reutilized during starvation. If fat oxidation is impaired the stored fat would be of little survival value. Thus the most important assumption behind the thrift family of hypotheses that the stored fat helps survival during periods of food crunch has not been demonstrated in humans (Speakman 2006).
viii. Is human physiology well adapted to feast and famine comparable to animals that hibernate or migrate over long distances? These animals store fat before hibernation or migration and utilize it through the non-feeding period with or without physical activity. In these animals, during the non-feeding period fat is preferentially utilized and until the last traces of fat are burnt out, proteins are not degraded. In humans muscle protein degradation begins much before complete depletion of fat stores (Pond 1998; Watve 2013). In brown bears that habitually hibernate, fat accumulation is not associated with adverse metabolic effects including insulin resistance, glucose intolerance and systemic inflammation (Arinell *et al.* 2012; Sommer *et al.* 2016; Stenvinkel *et al.* 2013). Black tailed godwits have a sufficiently plastic lipid metabolism that enables them to manage energy requirement of migration independent of diet composition (Viegas *et al.* 2017). On the contrary diet composition is believed to be a contributor to human obesity (Astrup & Brand-Miller 2012; Buettner *et al.* 2007; Miller *et al.* 1990; Tucker *et al.* 1997). It appears therefore that animal species evolved to adapt to a feast and famine challenge by storing and reutilizing fat, have also evolved mechanisms to avoid adverse metabolic effects of fat, if any. Human physiology does not seem to match that of animals evolved for thrift.
ix. The distribution and type of fat: Metabolically visceral versus subcutaneous fat and white versus brown fat are remarkably different and there is substantial individual variation in the distribution and type of fat (Avram et al 2005, Ibrahim 2009, Osama et al 2006, Saely et al 2012). While visceral fat is strongly positively associated with type 2 diabetes, atherosclerosis and other disorders, the subcutaneous and brown fat may even have protective effects. The thrift family of hypotheses does not account for the individual variance in distribution and type of fat.

Owing to the multiple flaws indicated in the thrift family of hypotheses there have been two different lines of attempts to explain human obesity. One is to propose refined or improvised versions of the thrift hypothesis that can escape one or more of the above criticism (Wells 2007). For example thrift is said to have evolved not under selection by harsh famines but by more subtle periodic changes in food availability (Prentice *et al.* 2008; Stipp 2011). The risk of starvation is argued to be sufficient to influence the strategy even when actual incidence of famine mortality is uncommon (Higginson *et al.* 2016). Interestingly some of the refinements of thrift concept refrain from mentioning thrift. For example the insurance hypothesis of Nettle et al (Nettle *et al.* 2017) is nothing but plastic and perception induced thrift. They argue that individuals store fat as a buffer against future starvation when the perceive food insecurity is high. But they neither mention the background of other thrift based hypotheses in their argument nor explain how their hypothesis differs from thrift. The refined versions of thrift try to overcome one or more of critical issues faced but none of them attempt to explain all of them. In addition, the insurance hypothesis suffers more problems as follows

1. Although the goals for global food security are still far from being achieved, there is a demonstrated increasing trend of food security in almost all parts of the world (Benson & Shekar 2006; Jamison *et al.* 2006). The rise in global food security is because of better agricultural productivity, quicker transport of food anywhere in the world, better storage conditions and greater social pressures on the governments to undertake nutrition and public health programs. The trend of increasing food security contradicts the trend of increasing obesity globally.
2. The epidemiological evidence given in support of the insurance hypothesis is weak since the correlation in meta-analysis is dominated by American women. Significant correlations are not found in men or children anywhere or women in low income countries. Further the American women data are not corrected for ethnicity. The ethnic representation in groups with different levels of food insecurity is likely to be different and whether the correlation remains significant after correcting for ethnicity remains to be examined.
3. Lastly a correlation between perceived food insecurity and mean body weight, which is the only epidemiological evidence in support, is possible because of reverse causation. Individuals with more uncontrolled hunger or food specific impulsivity are more likely to feel food insecurity. If only perceived insecurity is considered and not real life food availability, a reverse causation can lead to a significant correlation. The observed higher prevalence of food specific impulsivity in obese individuals (Houben *et al.* 2014) increases the plausibility of reverse causation.
4. An implicit assumption in the insurance hypothesis is that the predicted food crunch is not actually experienced. If it is experienced, the stored energy will get utilized lowering the adiposity. So the body weights are unlikely to be stably high. If food insecurity is perceived but not actually experienced, then escalation of adiposity is possible. This argument is in line with the central explanation of the obesity epidemic given by the thrift family of hypotheses. Perception of food insecurity without actual insecurity is needed for the rising obesity.

Therefore although food insecurity can potentially lead to increasing adiposity and the phenomenon may be theoretically sound and demonstrated in animal populations (Anselme *et al.* 2017); whether it is a predominant mechanism in the contemporary human population is still an open question.

The other reaction to the limitations of the thrift family of hypotheses is to find alternative explanations for human propensity for obesity and individual variation in it. Some notable examples of alternatives to thrift are drifty gene by Speakman (Speakman 2008) and behavioural switch by Watve and Yajnik (Watve & Yajnik 2007). We propose here an alternative explanation for the propensity for obesity based on behavioural optimization. The hypothesis is based on principles of foraging theory and suggests many testable predictions. We also discuss the implications of this hypothesis for the current global obesity epidemic and its control.

## The hypothesis

Species that experience predation, aggressive competition, extreme environments or any other foraging related risk, evolve mechanisms to optimize the trade-off between nutritional gains and foraging risk. Since the risk can be variable, there is plasticity based on perceived risk. The trade-off optimum depends upon pre-existing energy reserves and thus leads to a steady-state body weight. For such a species, if foraging risk is removed, the optimization mechanisms give way to overeating and escalated body weights. We propose that food intake regulation evolved in humans mainly under selection for foraging optimization. Feeding as well as stored energy induces lethargy and risk aversion which arrests foraging efforts and thereby feeding until energy reserves are utilized sufficiently. Since feeding is detached from foraging in modern life the regulation mechanisms do not work normally leading to a propensity for obesity. We explore this hypothesis at both ultimate and proximate levels.

## Foraging optimization

Foraging optimization has been an important focus of interest in behavioural ecology with substantial theoretical as well as empirical inputs. Animals have been modelled as well as demonstrated to optimize their foraging strategies to maximize nutrient gains (Clark & Mangel 1986; Kamil *et al.* 2012; Parker *et al.* 1990; Watve *et al.* 2016a, 2016b). Presence of a predator substantially alters behavioural and physiological patterns of a population (Preisser *et al.* 2005). Predation and other foraging risks necessitate certain trade-offs that affect total energy intake (Brown & Kotler 2004) as well as energy allocation (Higginson *et al.* 2012, 2016). Our approach is inspired by prior empirical as well as theoretical studies showing that foraging energy gains are traded off with predator risk and that can affect body weight as well as adiposity of an individual (Brown & Kotler 2004; Cresswell *et al.* 2009; Higginson *et al.* 2012, 2016; Lima 1986; Macleod *et al.* 2005; McNamara & Houston 1990; Zimmer *et al.* 2011). Models used so far invariably assume thrift directly or indirectly. The question whether the currently observed rise in obesity can be explained independent of thrift and can satisfactorily address issue i to xiii raised above remains unaddressed. Fat is treated by most authors only as a means of storing energy and several other functions of fat that can affect selection are not accounted for. There is an inevitable loss in energy due to thermodynamic and metabolic reasons, i.e. a very small part of the consumed energy can be stored in the form of fat and a further smaller fraction of the stored energy can be reutilized for metabolic purposes. This inevitable loss was also not explicitly considered by the earlier models. Further these models deal with an ecological optimum but do not incorporate the biochemical or neuronal mechanisms for achieving the optimum. None of the previous models explain the same phenomenon simultaneously at proximate and ultimate level which logical compatibility and synergy between the two levels.

We extend the logical framework of earlier models to formulate a new modelling framework that allows us to ask a set of new questions relevant to the origin of obesity and to make additional testable predictions. The new framework incorporates many subtle factors associated with fat ignored by the previous models and still remains mathematically simple enabling analytical solutions. In our formulation of the models foraging optimum and adiposity interact dynamically since stored energy influences the foraging optimum and optimum foraging decides how much energy is available for storage. We then proceed to apply these models to humans in an attempt to explain the recent obesity epidemic and make many testable predictions for our hypothesis.

## Body weight control by foraging regulation: Ultimate reasoning

For an individual the fitness benefits of food intake are assumed to be non-monotonic, there being an optimum food intake under a given set of conditions. Most foraging models use some immediate benefit of foraging such as energy gain. However in order to incorporate short term as well as long term cost benefits we use the overall contribution to lifetime fitness on the Y axis (Fig. 1).

**Figure 1:**
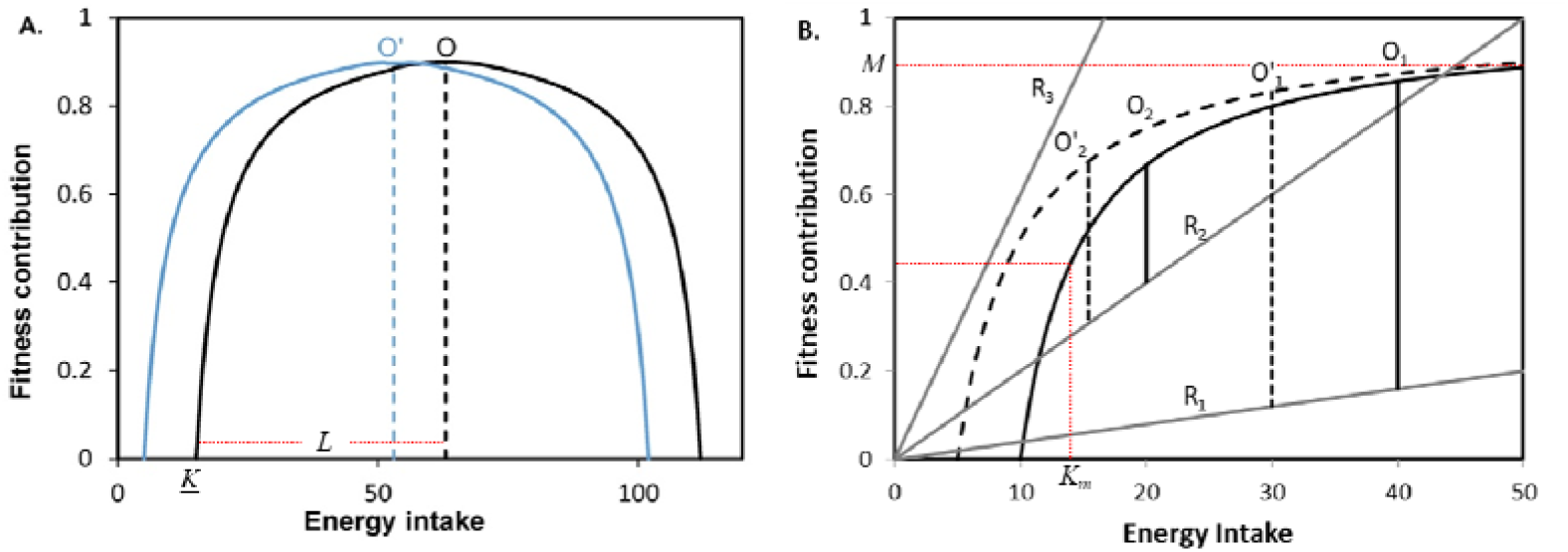
Fitness gains from foraging: (A) For species with little or no foraging risk (B) For species with varying degrees of foraging associated risks indicated by lines with slopes R1, R2 and R3. The foraging optimum lies where the difference between the gain and the risk is maximized. Note that the slope of the risk line influences the foraging optimum substantially. Optimum foraging for (B) is always more towards the left of that of (A) (note the difference in scale), i.e. the behavioural optimum is always lower than the metabolic optimum. Therefore in species that have consistently faced high foraging risk, metabolic optimization may not evolve or will remain weak. If this assumption is true we may use a saturation curve for (B) instead of an inverted U shaped curve. In both cases curves shift to the left if the individual has energy reserves. Accordingly the foraging optimum (now O’) also shifts.

We first assume an optimum food intake from metabolism point of view alone which could be approximated by an inverted U shaped curve the spread and height of which could be different for different species and different habitats. The inverse U shaped curve (Fig 1) need not start from zero as there might be a minimum energy intake K below which contribution to reproductive fitness is zero. After this X intercept the contribution to fitness increases to reach a maximum. This is the optimum food intake at which maximum fitness benefit is obtained. Above the optimum level food intake may cause metabolic problems or even physical problems such as greater energy expenditure required for locomotion. Since in wilderness food is obtained by foraging, we can say that there is an optimum level of foraging whose contribution to fitness is maximum. We assume here that the X axis of this curve represents net intake from foraging in which the cost in terms of energy and time spent in foraging is already subtracted. Y represents the fitness contribution from this intake. If an individual already has stored energy, the entire curve would shift to the left along with the optimum, proportionate to the amount of usable energy.

## Model without foraging risk

In the absence of any risk specifically associated with foraging, the optimum food intake lies where the curve reaches a peak. The two arms of the curve to the left and right of the peak need not be symmetric. We will assume them to be symmetric for simplicity of the model and then examine the possible effects of asymmetry. Assuming symmetry the inverted U shaped fitness curve can be represented by a simple equation

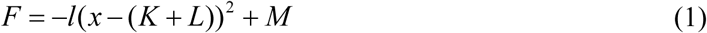

Where F is the fitness contribution of foraging

M is the maximum possible fitness which lies at the optimum food intake 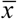, when 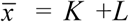 where K is the minimum foraging intake after which contribution to fitness begins. The constant *l* or *L* decides the shape of the inverted U.

*L* and *l* are not independent constants and *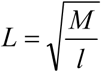* But we use two different symbols since they are associated with different intuitive interpretations. Whereas *l* decides the spread of the dome, *L* is the length from the X intercept to the optimum X. If an animal has stored energy, then *K* would reduce proportionate to the energy reserve such that the curve shifts to the left thereby reducing the optimum food intake *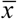*.

Apart from energy storage, adipose tissue has a number of other functions related to fitness that include metabolic, hormonal, immunological and reproductive functions (Pond 1998; Watve 2013). Therefore we assume that to a certain extent fat depot is a normal component of fitness functions even in the absence of selection for thrift. Therefore we first start with a simplistic assumption that fat deposited in the adipose tissue is directly proportional to the energy intake. This assumption is not critical to the model and we will show later that even if we assume some thrift, i.e. a pre-evolved tendency to store fat when food is available, the model and its central result remains unchanged. A fraction of stored fat is burnt per unit time equal to a constant basic metabolic rate *B*. Alternatives to both these assumptions are considered later in sensitivity analysis.

Thus adiposity at time *t +1*

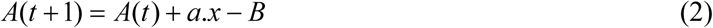

Where *a* is the energy deposited in adipose tissue per unit energy intake and *B* is the basic metabolic rate.

Adipose tissue can reduce the minimum foraging efforts *K* such that

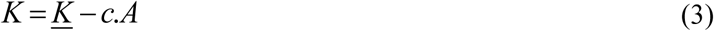

Where, *c* is a constant for equivalence of utilizable stored energy per unit foraging energy intake. Due to thermodynamic considerations, *c* << 1. *<u>K</u>* is the minimum amount of energy intake needed for getting fitness benefits in the absence of any stored energy.

Thus there is a dynamic relationship between equations 1, 2 and 3. The optimum 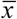 given by equation 1 affects *A, A* alters *K* and *K* decides 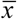. A steady state in this dynamics is possible when

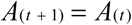

This can happen when *a. 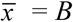.*

It can be seen that this equilibrium is stable because when *A* is small, *K* becomes large and thereby 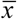 increases adding to the fat depot. When *A* is large, 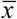 decreases and thereby *A*. The steady state 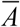 is given by

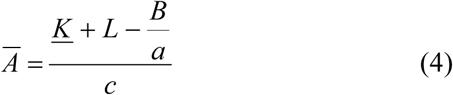

It can be seen now that there is a steady state body weight provided there is a proximate mechanism to optimize foraging to maximize the gains. The symmetry or asymmetry of the inverted U shaped curve does not affect the steady state for the baseline model. It can however influence the evolution of thrift, which can be defined here as foraging more than the optimum to store additional fat. This will invariably lead to a fitness loss but if the decline on the right hand side is not steep then under the threat of somewhat predictable food shortage, thrift can evolve. We can say that if there is evolved thrift, the actual food intake is somewhat higher than the optimum i.e. *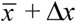.* There will be an optimum Δ *x* depending upon the perceived or actual food scarcity. However since the optimum shifts to the left on having stored energy, there will be an upper limit on body weight independent of whether the period of food scarcity is actually faced or not. If there is evolved thrift of Δ *x* and no food restriction faced as predicted, the steady state body weight will be

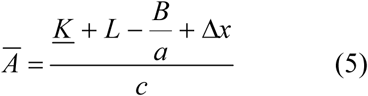

This implies that evolving in feast and famine situation but not facing a famine in modern life is not a sound reason for escalating body weight. The adiposity will remain at a steady state as in eq. 5 even if no food crunch is ever experienced. Further if this level of adiposity is repeatedly reached in the evolutionary history, we expect mechanisms to evolve that would prevent adverse metabolic effects of the evolved level of obesity as seen in periodically hibernating animals. Adverse metabolic effects are likely to be seen if a level of adiposity is achieved which is not repeatedly faced during evolution. This is unlikely to happen by an evolved mechanism from thrift.

## Model with foraging risk

The optimum changes substantially if there is a risk associated with foraging. The risk can come from predators, aggressive competitors or harsh environments such as extreme temperatures. The total risk is likely to increase in direct proportion to the time spent in foraging, which is represented by a straight line with positive slope (Fig 1 B). Optimum energy intake in this model is where the difference between the fitness curve and risk line is maximized.

It can be seen that the risk optimum is always towards the left of the metabolic optimum (Fig 1 B). This implies that animals that have faced high foraging risk for several generations may fail to evolve metabolic regulation or may weaken or lose metabolic regulation mechanisms since they would always remain unused. For such species the optimization model may be based on a saturation curve rather than on an inverted U shaped curve (Fig 1 B). It is possible therefore that in some species the metabolic regulation mechanisms predominate and in other species the behavioural regulation mechanisms evolved for risk trade-off predominate. Species in which the risk level at resting and foraging is not substantially different are expected to be metabolic regulation species and the ones facing high level of foraging risk as compared to resting risk are likely to evolve as behavioural regulation species. Animals such as elephants that have little foraging associated risk are likely to be metabolic regulation species. Large ground herbivores that have similar predator risk during resting as well as foraging will also constitute metabolic regulation species. Species that have predators but also have safe refuges for resting as well as species that face strong aggressive competition during feeding will belong to behavioural regulation species. The foraging and thereby food intake optimum for the two categories is expected to be different being at the highest point of the inverted U (Fig 1 A) for metabolic regulation species and much towards the left of metabolic optimum for behaviour regulated species.

The model with foraging risk assumes that individuals have a safe resting place and foraging risk increases linearly with foraging effort. There is a minimum energy intake *K* required above which further energy intake contributes positively to fitness. For *x > K*, the fitness benefit *F* obtained from net foraging energy gain *x* is given by

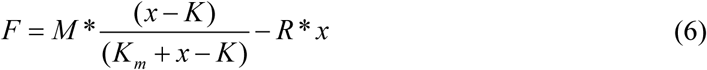

Where *F* is fitness benefit from foraging

*M* is maximum fitness benefit obtainable from a foraging bout

*R* is the slope of the risk line

*K* is minimum energy required for positive fitness returns

*X* is net energy gain from foraging

*K*_*m*_ is the half saturation constant for the benefit curve.

From equation 6 we can write

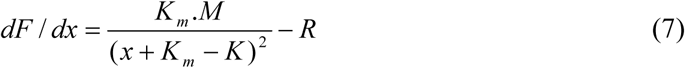

Fitness is maximized when *dF/dx =0*, which can be achieved when

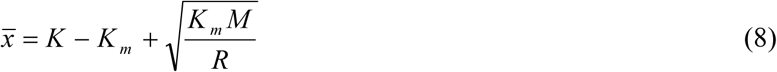

Where, 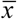 is the optimum foraging energy intake.

With this optimum 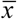 and the steady state considerations as above (eq. 2 and 3) steady state adiposity is obtained as,

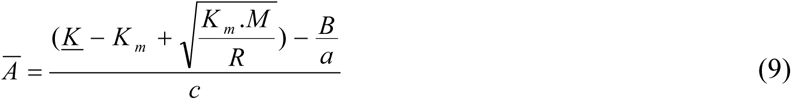

Thus according to the model steady state adiposity is a non-linear decreasing function of foraging risk *R* (Fig 2a).

**Figure 2:**
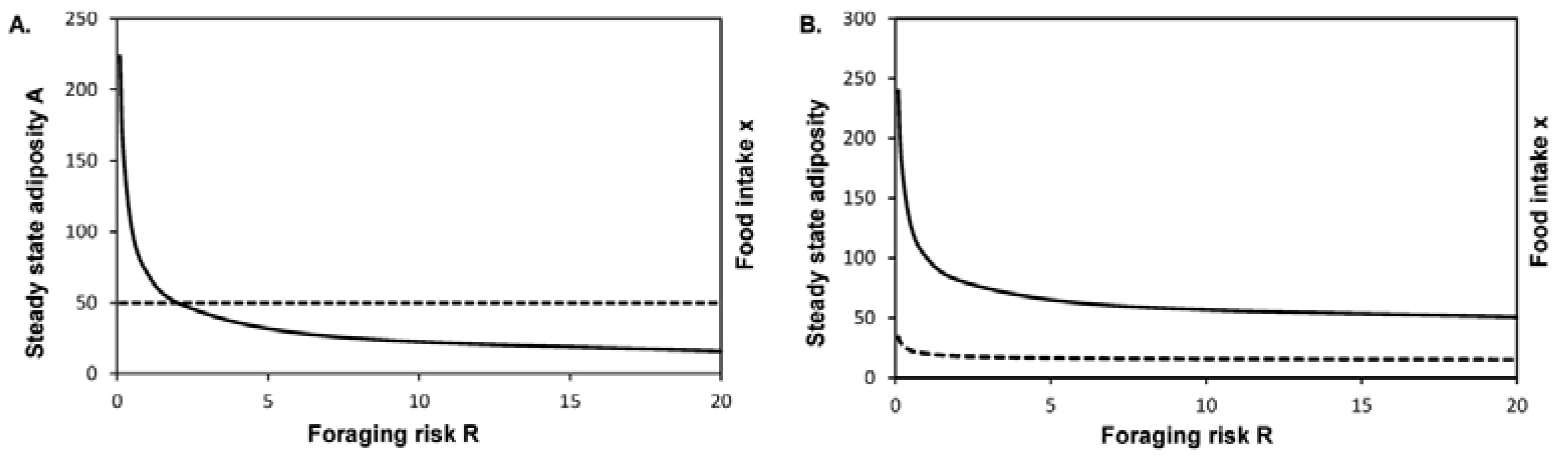
Steady state adiposity at different levels of foraging risk: Other parameters being constant (*M* = 1000, *L* = 5, *<u>K</u>* = 55, *a* = 0.1, *B* = 1, *c* = 1 and *R* is varied between 0.1 and 20) it can be seen that if the risk level reduced below a threshold, steady state adiposity (solid line) increases dramatically but the food intake (dotted line) may not increase proportionately. A) Assuming resting energy expenditure to be constant, adiposity is uncorrelated to food intake B) Assuming resting energy expenditure a function of body mass, the food intake may increase along with adiposity at low risk levels.

Over a wide range of foraging risks the curve remains relatively flat, i.e. there is little increase in adiposity per unit risk reduction. However if the risk reduces below a threshold, the steady state adiposity rises almost like a wall. If in the modern urban lifestyle the risk levels have fallen below the threshold, then a level of adiposity is expected which was never faced in the evolutionary history. As a result the physiology is most unlikely to be adapted to this level of obesity and therefore adverse metabolic and endocrine effects may be seen.

Interestingly in this baseline model, the steady state food intake does not increase with body weight at lower risk levels although longitudinally food intake will be higher during weight increase. The steady state adiposity will be affected by *B, a, <u>K</u>*, *K*_*m*_ and *c*. However there is no a priori reason why these parameter would change dramatically with lifestyle. They are evolved physiological constants that can have a limited individual variation. On the other hand foraging related risk is highly variable in natural ecosystems according to time of the day, season, predator movements and other cyclic or stochastic variables. Therefore a plastic response based on perceived risk is expected to evolve. A dramatic decrease or almost absence of feeding related risk is certainly a component of modern lifestyle which alone is sufficient to escalate adiposity. This can happen independent of thrift or any other mechanism specifically evolved for fluctuating food availability. Therefore it is the most likely causative factor in escalation of adiposity.

Similar to the model without risk, evolution of thrift can interact with foraging optimization with risk, but the interaction can be two way. Since the risk levels can change substantially to forage more when perceived risk is low and to avoid foraging when it is high is a good strategy. This will happen naturally through the optimization program and no separate evolved mechanism of thrift is needed. On the other hand if thrift has already evolved for fluctuations in food availability, it can increase the escalating effect of lifting foraging risk. Again similar to the arguments above, absence of famine alone is not a sufficient reason for escalation of adiposity. Even if a famine is never faced adiposity will remain stable at

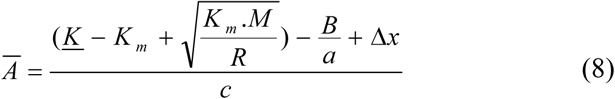

On the other hand if *R* approaches zero, evolved thrift of Δ *x* can escalate adiposity further.

The model requires that mechanisms of sensing the current foraging risk should have evolved. They include predator sensing by smell, conspecific and heterospecific alarm calls, visual signs and experiential knowledge about dangerous times, places and seasons. As a result the foraging optimum would be plastic and will depend upon the perceived risk at a given time.

In presence of foraging risk, it is adaptive to avoid foraging when stored energy is sufficiently high. Therefore any mechanism of avoiding foraging drive mediated by adiposity would be adaptive. This may include feeding induced or obesity induced lethargy and risk aversion tendency. We suggest that lethargy is not an inevitable consequence of overfeeding or obesity but is an evolved mechanism to optimize foraging efforts. Not all species show signs of lethargy upon feeding to satiation (Nie *et al.* 2017). But for species with large foraging specific risk, feeding induced and/or obesity induced lethargy would be specifically adaptive. In the absence of such specific adaptive role of lethargy, it would be absurd to believe that having extra energy makes an individual lethargic.

## Sensitivity analysis

We will now relax or alter the assumption of the baseline model to see whether the main result, that is risk reduction is sufficient for increase in adiposity, is qualitatively sensitive to any of the challengeable assumptions of the model.

1. We have assumed that the basal metabolic rate *B* is constant. Many studies show that it increases with body weight (Hall 2007; Leibel *et al.* 1995; Lissner *et al.* 1990). We altered equation 4 to incorporate this as

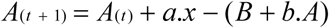

Where, *B* is the basal metabolic rate of lean tissue and *b* the metabolic rate of unit adipose tissue. After incorporating this change in the assumption, the relationship of *A* with *R* remained qualitatively similar but the steady state food intake increased with adiposity (Fig 2 B). This was because individuals with higher adiposity needed to take in more energy in order to compensate for the increased metabolic consumption. In humans, adiposity is associated with decreased physical activity (Must & Tybor 2005). It is therefore likely that the net energy consumption may increase, decrease or remain the same with increasing adiposity. Since in a steady state the energy intake and expenditure is equal, there can be a positive, negative or no correlation between adiposity and energy intake. An important epidemiological implication of this model is that BMI or any other measure of obesity need not be positively correlated with energy intake in a steady state. There may be no correlation or even negative correlation.
2. Distance between refuge and foraging grounds: If we assume that the safe resting refuge is at a substantial distance from the feeding grounds then there would be a travelling cost associated with every foraging bout. In such cases the overhead cost of foraging will increase. However Watve *et al.* (2016a) show that in models that optimize the benefit cost difference, overheads do not change the position of the optimum in relation to the curve. Therefore even in this set of assumptions the relationship of *A* with *R* remains unchanged.
3. Norms for fat deposition: We have assumed in the baseline model that fat deposited is directly proportional to the net energy intake. We can alter this assumption. It is possible that energy is preferentially driven to reproduction and secondarily to storage. In that case we may write

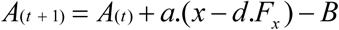

Where *F*_*x*_ is the fitness possible at a given *x* and *d* is the energy requirement for fitness related activity. This means that when allocating more energy to reproductive efforts does not yield proportionately more fitness, the excess energy gets stored. Alternatively the energy obtained may be first utilized to meet the basal metabolic demand and a proportion of the excess is stored as fat. Thus

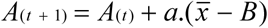

Analytical solutions with one or combinations of these assumptions do not change the nature of the relationship between *A* and *R*. Thus the main result of the model is robust to the array of alternative assumptions about energy storage.
4. Everyone does not have to forage: In human societies there is division of labour and food sharing so that every individual need not forage for feeding. If we assume that one individual forages for an entire family, the optimization of the non-foraging individuals may be different. However, closer examination revels that in a typical human family structure, loss of or injury to a foraging individual is a loss to the non-foraging members of the family as well. Therefore the foraging risk trade-off operates in a qualitatively similar manner.

In summary, in the foraging under risk model, as long as the foraging benefit follows a saturation curve, steady state adiposity follows a non-linear relationship with foraging risk such that for a large variation in foraging risk, little change is observed in the adiposity. However if the risk reduces below a threshold there is a sudden rise in mean adiposity. This result is robust to many variations of the model.

## Biochemical mechanism of obesity control: Proximate reasoning

If there is an optimum level of foraging, the foraging individuals need to evolve mechanisms to achieve the optimum. The mechanism needs to be sufficiently plastic to accommodate the variable levels of food abundance and risk levels in a given environment. Such mechanisms have been described and are twofold involving and not involving risk perception. The risk independent foraging regulation is a likely result of feeding or adiposity induced lethargy. The model assumes the optimum to shift to the left on having energy reserves. At a proximate level feeding or adiposity induced lethargy can bring about the leftward shift in foraging effort. A number of long known classical biochemical mechanisms are implicated in food induced lethargy. Glucose, insulin and cholesterol upregulate serotonin expression (Fernstrom & Wurtman 1971; Golomb 1998; MacKenzie & Trulson 1978). Fatty acids compete with tryptophan for albumin binding (Bohney & Feldhoff 1992; Cunningham *et al.* 1975) and the free tryptophan can cross the blood brain barrier to facilitate serotonin synthesis. Serotonin, in turn, induces central fatigue, drowsiness and lethargy (Cotel *et al.* 2013; Soares *et al.* 2007). Adipose tissue secretes a number of pro-inflammatory cytokines which also contribute to a fatigue and lethargy response (Norheim *et al.* 2011). The specific molecular pathways to lethargy and absence of feeding induced lethargy in some species (Nie *et al.* 2017) suggest that lethargy could have specifically evolved for foraging regulation. Such a regulation is bound to fail when feeding is detached from foraging.

The other class of mechanisms of foraging regulation work through risk perception. A number of fear signals in the brain are mediated by perceived risk, aggression and anxiety. Many of the peptides expressed by fear have anorectic action. The list of regulators inhibiting food intake is considerably longer than that of appetite stimulators (Watve 2013) and many of the peptides inhibiting food intake are also the ones mediating fear reactions, whereas the majority of the agents reducing anxiety responses stimulate appetite (Adam & Epel 2007; Bazhan & Zelena 2013; Kask *et al.* 2000; Smith *et al.* 2006). Of particular interest are the brain signal molecules, α melanoctye stimulating hormone (α-MSH) (File 1981; Kokare *et al.* 2005), histamine (Hasenöhrl *et al.* n.d.; Mercer *et al.* 1994; Ookuma *et al.* 1989; Privou *et al.* 1998), acetylcholine (Herman *et al.* 2016; Stipp 2011) and cocaine and amphetamine regulated transcript (CART) (Asakawa *et al.* 2001; Kask *et al.* 2000; Stanek 2006), all of them being related to fear, anxiety and risk avoidance on the one hand and decreased food intake on the other. It is very likely therefore that signalling by peptides like CART and histamine in the brain has evolved to fine tune the trade-off between energy gain by foraging and associated risk. When a higher risk is perceived, higher levels of these signals are generated which will reduce hunger, the main foraging drive, and thereby help in achieving the optimum in nutrition-risk trade-off. In the absence of both predators and competitors, rats deplete their CART levels and become hyperphagic and gain weight (Nakhate *et al.* 2011; Upadhya *et al.* 2013). Appetite control is a complex interplay between short term metabolic signals such as insulin, long term signals such as leptin and behavioural signals such as CART and alpha MSH. The action of leptin is at least partially CART dependent and CART levels are fine-tuned by perceived fear and aggression (Elias *et al.* 1998; Elmquist *et al.* 1999; Kristensen *et al.* 1998). It has been shown that the difference between diet induced obese rats and diet resistant rats (those that fail to become obese when kept on a high fat diet) lies in the CART and alpha MSH levels in the hypothalamus (Bouret *et al.* 2008; Tian *et al.* 2004). If CART and alpha MSH levels are low, food intake regulation fails and animals become obese. This may be the key to behaviour induced suppression of appetite. On the other hand if an animal is starving, or if leptin signalling is low, CART expression is suppressed (Kristensen *et al.* 1998; Tian *et al.* 2004; Yang *et al.* 2005). This would impair the fear response and the animal will be prepared to take greater risk for foraging when desperate for food.

It is unlikely to be a coincidence that molecules involved in anxiety are also involved in appetite control. Leptin target neurons in the arcuate nucleus of the hypothalamus include POMC (Proopiomelanocortin) -expressing neurons. These produce and secrete various signals from axon terminals, including α-MSH, β-endorphin and CART (Cowley *et al.* 2001; Kristensen *et al.* 1998; Millington 2007). Combined, these neurotransmitters and neuropeptides are the candidate effector molecules for mediating energy homeostasis and glycemic control (Balthasar *et al.* 2004; Claret *et al.* 2007; Cone 1999; Sainsbury *et al.* 2002; Schwartz *et al.* 2000; Woods *et al.* 1998). The leptin, CART, alpha MSH interaction appears to have evolved in such a way to achieve an optimum trade-off between the metabolic demands of the body and the risk associated with foraging for food. We will model this interaction now taking CART as a representative of all fear peptides that interact with leptin to exert a regulation on hunger response.

**Figure 3:**
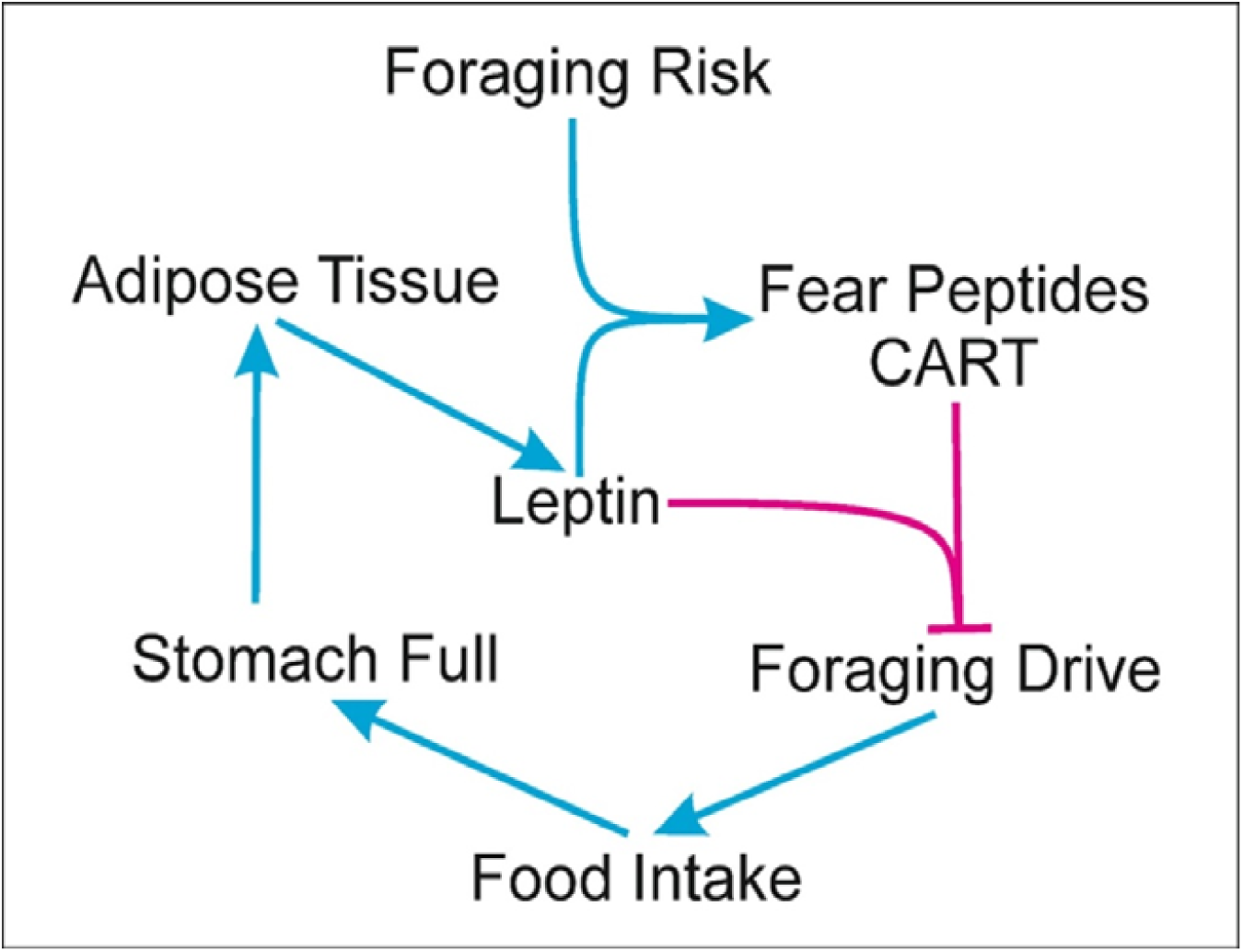
Schematic representation of Leptin-CART interaction in hunger control: Blue arrows denote facilitatory and red blunted lines inhibitory effects. A model for this interaction gives a relationship between foraging risk and adiposity similar to the foraging optimization model at behavioural level.

Feeding and foraging drive, i.e. hunger, is modulated by leptin levels as well as CART levels. A simple way to model it is

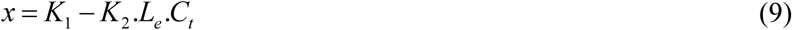

Where *L*_*e*_ represents the level of leptin and *C*_*t*_ the level of CART in the brain, *K*_1_ is the maximum capacity to feed at a time and *K*_2_, a rate constant for the joint action of *L*_*e*_ and *C*_*t*_, we continue to use *x* in the same sense as the foraging model above.

Since leptin is produced by the adipose tissue

*L*_*e*_ = *s.A* and since CART expression is positively affected by leptin

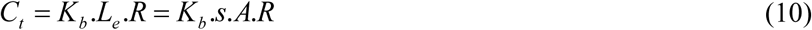

Therefore

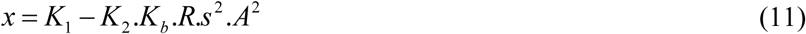

Considering equation 11 along with equation 2, steady state adiposity can be written as

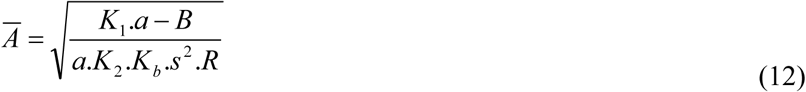

In this equation, parallel to the foraging optimization equation, the steady state adiposity is inversely proportional to the square root of risk and as a result below a threshold risk the steady state adiposity increases sharply. Thus the relationship between steady state adiposity and perceived risk remains the same in the proximate and ultimate model. Here again, at steady state the food intake need not increase in proportion to adiposity. Convergence of an optimization model of behavioural ecology and a model of demonstrated molecular interaction makes the joint proximate-ultimate model more promising for understanding the root causes of obesity. On the other hand no definite biochemical mechanism for thrift has been convincingly demonstrated. Therefore we do not have a proximate model of thrift to compare with the ultimate model.

## Sensitivity analysis

We will now relax or modify the assumptions of the proximate model to see whether the relationship between *A* and *R* changes qualitatively. We assumed the anorectic action of leptin to be linear and directly proportional to CART levels in the brain. It might be more appropriate to consider non-linear functions as in the equations below.

Foraging drive or foraging modulated non-linearly by leptin levels:

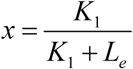

Since brain cart levels (C) mediate the action of leptin, which also need not be linear,

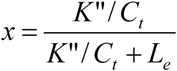

The effect of leptin on the expression of *C*_*t*_ can also be considered non-linear and increase with a saturation relationship so that

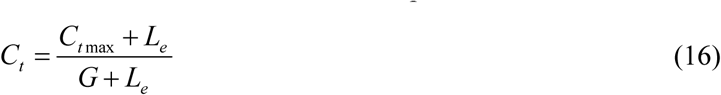

We also challenge the assumption of basic metabolic rate being constant and make it a function of *A* as in the foraging model. The non-linear models make analytical solutions more complex but the relationship between steady state adiposity and perceived risk remains unaltered.

This demonstrates the robustness of the *A* and *R* relationship. It also demonstrates the convergence of the ultimate and proximate models.

At a biochemical level the mechanisms that we modelled are already well demonstrated. Not only predator presence but even extreme temperatures trigger CART expression (Lau *et al.* 2016; Sánchez *et al.* 2007; Song *et al.* 2012) suggesting that the mechanism might cover different types of foraging risks. Apart from the leptin-CART interaction that we formally modelled here, there are other supplementary mechanisms that would help the trade-off. Apart from leptin, high energy diet leads to acute upregulation of hypothalamic CART (Archer *et al.* 2005) which will have a short term role in foraging regulation. Further the neuro-behavioural effects of CART are different in starved versus well fed animals (Davidowa *et al.* 2005; Yang *et al.* 2005). CART also interacts with other hunger related signals including NPY and cholecystokinin (Heldsinger *et al.* 2012; Lambert *et al.* 1998) in a similar way.

In addition to leptin and CART, there are other mechanisms that can mediate the interaction between hunger related signals and fear, risk or aggression related signals. Serotonin suppresses aggression (Ferris 1996; Ferris *et al.* 1997), increases fear and anxiety (Frick *et al.* 2015) and increases central fatigue, lethargy and drowsiness (Davis *et al.* 2000; Meeusen *et al.* 2006). Serotonin is also a satiety signal and arrests feeding response (Simansky 1995; Voigt & Fink 2015). Common mechanism for satiety and aggression suppression would be important for a species with high food related aggression. When predators and scavengers fight over a kill, there is a high risk of injury. When an animal is desperate for food it has low serotonin and would be aggressive, take risk and feed on the kill. However having gobbled some food, the serotonin levels will start rising and the individual would tend to avoid aggression and retreat from the fight. Such a mechanism would keep food intake in check in presence of other aggressive competitors but will fail in their absence. Food intake is shown to increase corticosteroid expression (Al-Damluji *et al.* 1987; Alexander *et al.* 1995; Gibson *et al.* 1999; Laugero 2001; Sander 1992) which in turn reduces risk tolerance (Thaker *et al.* 2009). All such mechanisms are compatible with the foundation of the model that satiety or large energy reserves should suppress physical activity, aggression and risk tolerance and reciprocally high risk perception should suppress hunger and physical activity. The multiplicity of mechanisms indicates that there could be strong selection pressure for mechanisms of optimizing nutritional gains versus foraging risk.

Apart from the net energy dynamics, energy allocation to lean tissue, particularly muscle versus fat is also likely to be affected by predatory pressure (Higginson *et al.* 2012). Greater muscle activity is needed for an active anti-predator response whereas higher fat is likely to increase predator susceptibility. Therefore predatory pressure should increase muscle mass and decrease fat mass. Here again CART-leptin interaction has a possible role to play. CART^−/−^ mice had reduced lean mass and increased adiposity (Asnicar *et al.* 2001) indicating that this system plays a role in energy allocation too. Further catecholamine and CART modulate browning of fat and thermogenesis (Reddy *et al.* 2014; Richard & Boisvert 2010). Thus a large number of known biochemical mechanisms are compatible with the foraging risk optimization model and are they unlikely to find any alternative adaptive explanation.

## Comparison with earlier foraging models

Although foraging theory has been used earlier for explaining body weight regulation in the context of origins of obesity (Brown & Kotler 2004; Higginson *et al.* 2012, 2016), our model includes some realistic parameters and processes not explicitly included by them. We ask the question, for the first time, whether obesity is possible independent of thrift. This is a first attempt to clearly distinguish between thrift and non-thrift explanations of obesity. In spite of including many realistic conditions for the first time, the mathematical framework of our model is simple enough to allow analytical solutions. Although the model begins with a set of simplistic assumptions we show that it is quite robust to alterations in the challengeable assumptions.

One characteristic feature of human obesity is the apparent constancy of weight for an individual. Some of the earlier models explained it assuming and including a set point in the model in some or the other way (Higginson *et al.* 2016). Our model could explain a steady state weight for an individual without having to specifically incorporate a set point in the model. The steady state weight in the biochemical model is decided by CART levels and parameters of CART-leptin interaction. This notion is testable in principle, although it is difficult to get data on levels of CART and other peptides in human brain.

Our model explicitly includes thermodynamic and metabolic loss of energy during bioconversions. All known biochemical pathways have substantial loss of free energy. Therefore converting ingested food into stored fat during feast and reutilizing it again is a highly inefficient process and such a mechanism would evolve only when profitable in spite of the big loss of energy. The thermodynamic loss needs to be explicitly incorporated in any model of thrift.

Freedom from predation has been proposed to be responsible for rising obesity by Speakman (2006). However this model needs genetic drift over a period sufficiently long to for spread of obesognic mutants in the population under relaxed selection. Although freedom from predation is the common consideration in the two models, our model works at a physiological level and does not depend upon genetic drift. On the other hand freedom from predation may not be sufficient to increase obesity in the population by our model. Other foraging risks need to be eliminated simultaneously.

## The origin of human obesity

We recognized earlier a dichotomy of metabolic regulation versus behavioural regulation of food intake. We expect humans to be a behavioural regulation species more than a metabolic regulation species. Through much part of human history our ancestors were mainly scavengers competing with other large carnivores and aggressive scavengers. This is known to involve intense aggression on a kill being equivalent to a large value of *R* in the model. In addition the food distribution being patchy there is likely to be intra-species competition and occasional fights over food. The patchiness of food distribution would also apply to fruit and other plant food sources palatable to humans. Human ancestors also were exposed to predation until recently (Watve 1993). Therefore it is likely that behaviour regulation mechanisms for energy intake evolved to be stronger in humans and metabolic regulation might have remained weak if not absent. The origin of the prevalent human obesity pandemic according to our model is that in the modern lifestyle feeding is uncoupled from foraging and/or foraging is uncoupled from risk. When feeding is foraging dependent, feeding induced or fat induced lethargy is sufficient to arrest further food intake. However, if food is available without physical efforts, this control is bound to fail. Further if there is little risk associated with feeding, the leptin CART interaction will be impaired. When the evolved biological mechanisms for food intake regulation fail to work and we have to rely on cultural and cognitive restrictions on eating.

It should be realized from the model that a large change in foraging risk or levels of CART and other fear induced signals is tolerated without much change in body weight. Only below a threshold of these signals there is a rapid rise in adiposity. It is likely that human lifestyle started changing in the late stone-age starting with burial practices which might have reduced predatory pressure (Watve 1993). Subsequently food storage, animal husbandry and agriculture reduced the reliance on foraging for every feeding. Still as long as there were sufficient environmental triggers for expression of CART and other signals, the evolved signalling mechanisms for energy regulation could have worked efficiently. Protecting the animal herds from natural predators as well as from other people, protecting the crops from wild herbivores, protecting the stored food from conspecific and heterospecific robbers required some physical aggression and risk taking which would trigger the molecular signals. Therefore it is likely that until recently humans had sufficient levels of these signal molecules in the brain that would maintain a healthy control over food intake. With modern lifestyle all such signals have considerably weakened. Extreme heat and cold also poses foraging risk and is known to trigger CART expression. Pigs exposed to heat stress attain less weight in spite of food availability (Hicks *et al.* 1998; McGlone *et al.* 1987; Rauw *et al.* 2017). Exposure to ambient temperature variability is also substantially reduced in the modern urban lifestyle and temperature regulating devices is perceived as a plausible cause of the obesity epidemic (Johnson *et al.* 2011; Keith *et al.* 2006). Chronically reduced levels of CART and other risk signals is a promising model of explaining the increasing obesity.

## Explanatory power, evidence and testable predictions of the model

1. Why everyone is not obese: If there are individual differences in the perception of risk or the mechanisms by which risk suppresses foraging, there would be individual differences in resultant adiposity. It can be seen (Fig 4) easily that in the flatter portion of the curve, with some variability in risk perception the resultant variability in body weight is small but below the risk threshold the slope becomes very steep and as a result for the same variability in X axis, the variability in adiposity can be very large. Similarly in the proximate model, substantial variation in the leptin CART interaction parameters can be tolerated with little change in adiposity in the higher range of CART levels but at low CART expression levels the same amount of variability in the parameters of CART-leptin interaction can result into large differences in adiposity. Further in the flatter right hand side of the curve, where the slope is nearly constant, a Gaussian distribution on X axis will result into a Gaussian distribution on Y axis. However at low CART levels, since the slope changes rapidly, a Gaussian distribution on the X axis is expected to lead to a Y distribution with a positive skew. Thus we expect that along with an increase in mean adiposity there will be an increase in variance and a positive skew in the distribution. Data on Swiss conscripts (Bender N et al, manuscript under review) show that over the last century not only the mean BMI increased but the variance increased along with a positive skew. This change is compatible with our model.

**Figure 4:**
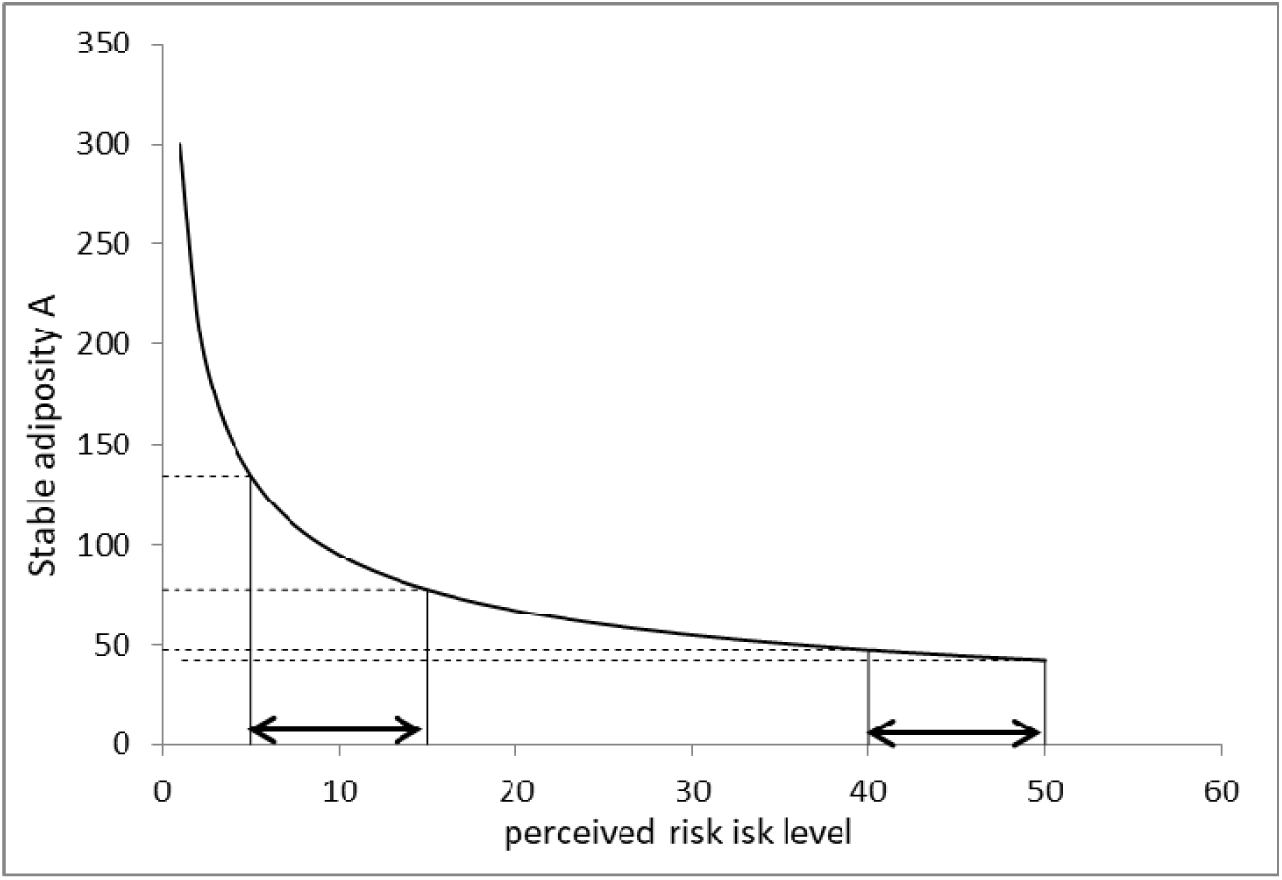
Effect of foraging risk perception on steady state adiposity: At a given risk level faced by a population individual variation in risk perception or CART-leptin interaction parameters results into variance in stable body mass. For equal variance on the X axis the variance on Y axis can be substantially different in different parts of the curve. At low risk or CART levels, both mean as well as variance in body mass is expected to increase.

With the beginning of agricultural and urban society, the foraging related regulation started becoming weaker. Perhaps in response to that different cultures appear to have developed different norms, customs and taboos on eating behaviour such as fasting, seasonal taboos on certain types of food, restrictions on time of eating or sequence of eating such as elderly first or kids first etc. Cognitive and cultural regulation of eating behaviour is likely to have increased its relative importance as foraging became gradually irrelevant. In the modern society awareness about fitness, diet and obesity is increasing and establishing up a new set of norms. Such norms have gained importance since the evolved regulation mechanisms became increasingly non-functional. Here again individuals differ in their compliance to such norms. Therefore both biological and cognitive differences are likely to be contributing to individual differences in the propensity to be obese.
2. Compatibility with foetal programming: The foraging hypothesis has no element which contradicts the foetal programming or developmental plasticity hypothesis. Individuals with intra-uterine growth retardation (IUGR) are more likely to accumulate fat if energy dense food is accessible in later life (Heerwagen *et al.* 2010; Langley-Evans *et al.* 2005; Malo *et al.* 2012; Prentice 2003). The association between low birth weights or other markers of IUGR and obesity, diabetes and CVD is observed over a wide range of populations (Boney *et al.* 2005; Curhan *et al.* 1996; Hirschler *et al.* 2008; Prentice 2003). This association is variously interpreted as thrifty programming in prediction of future energy limitations or a marker of food insecurity (Nettle *et al.* 2017). An alternative interpretation is that developmental constraints may limit physical strength of an individual who is more likely to live a socially subordinate life in a typical primate society. The social as well as ecological behaviour of a subordinate individual is significantly different than a strong and dominant individual. In food related competition, they are likely to be less aggressive and more opportunistic feeders. Such an individual will have a more leftward foraging optimum. In order to compensate for this loss they are likely to be programmed to spend less energy and reserve more. There is a developmental constraint on muscle mass since muscle cells do not increase in number in adult life. Further being aggression avoiders, they can also afford to invest less in muscle. This would increase *a* in our models leading to higher adiposity. Thus the risk optimization model is not only compatible with IUGR hypothesis but supportive of it.
3. Why impairment of fat oxidation marks obesity: Risk signals like CART also plays a role in energy expenditure (del Giudice *et al.* 2001; Kong *et al.* 2003). This makes sense because it is a high risk and therefore high CART expression context that foraging should be avoided and stored fat should be mobilized. The sympathetic nervous system which also reflects perceived risk is an important signal for fat mobilization (Bartness *et al.* 2010; Weiss & Maickel 1968; Yoshimatsu *et al.* 2002). This suggests a possible explanation for the observed impairment of fat oxidation seen in obese humans. The appropriate context for fat utilization is when foraging risk is high and foraging should be avoided. In such circumstances survival on stored energy is necessary. Therefore a high risk perception should trigger fat mobilization. In the modern urban lifestyle the deficiency of foraging and related risk creates a deficiency of appropriate signals for fat mobilization resulting into the observed impairment of fat oxidation mechanisms associated with obesity.
4. Addressing critical issues i to viii: We started by reviewing the issues contradicting the thrift family of hypotheses. It can be seen that the foraging optimization hypothesis is not faced with any of these issues. Our hypothesis does not assume any feast and famine selection, although it is compatible with it. The only assumption about human ancestry is an inevitable link between feeding, foraging and risk. It does not require genetic polymorphism to explain the population variance in the propensity to be obese (see below). Since the origin is not mainly genetic, GWAS is not expected to explain major part of population variance. The hypothesis is compatible with foetal programming but is not dependent on it which explains why foetal programming explains only a fraction of the epidemiological pattern. Further our hypothesis is also compatible with the behavioural switch, soldier diplomat dichotomy or behavioural deficiency hypothesis (Belsare *et al.* 2010; Watve 2013; Watve & Yajnik 2007) which explains multisystem involvement. Finally unlike thrift family of hypotheses, our hypothesis has well demonstrated proximate mechanisms for execution of the optimization and conditions for the failure of regulation.
5. What decides fat distribution: Visceral versus subcutaneous fat have different metabolic and endocrine effects, however most hypotheses about the evolutionary origins of obesity do not consider the proximate and ultimate causes of differential fat distribution. A moderate level of subcutaneous fat can be helpful in a risk and aggression prone lifestyle by cushioning and partially protecting against impacts and attacks. Therefore on facing risk it would be a good strategy to preferentially burn visceral fat and reserve subcutaneous one in anticipation of attacks, fights or accidents. Compatible with this is the fact that subcutaneous and visceral fat are differentially innervated and neuronal inputs are important regulators of fat metabolism (Bartness & Song 2007; Kreier *et al.* 2002). Absence of aggression and risk perception is therefore expected to increase visceral fat more than subcutaneous fat. Further since excess fat can be detrimental in presence of a predator, it would be adaptive to burn off more reserves rather than store. The physiological mechanism of achieving this is that CART and catecholamine signalling is an important driver of fat browning (Reddy *et al.* 2014; Richard & Boisvert 2010) absence of which may increase the proportion of white adipose tissue. Thus the signalling mechanisms involved in foraging regulation are also involved in deciding fat distribution and adipose tissue type. It should be noted that the thrift family of hypotheses have not offered any explanation for the type and distribution of fat at proximate and ultimate levels.
6. Food intake, energy reserves and lethargy: According to our model, developing lethargy after a heavy meal or on having high energy reserves is an evolved adaptive strategy. This is expected to evolve only in species that have a substantial difference in the resting versus foraging risk. Species that are exposed to predation risk even during resting should not evolve feeding induced lethargy. Although post-feeding lethargy is rarely studied quantitatively in any species, in at least one species of fish, physical activity, critical swimming speed and fast-start escape speed was shown to be unaffected by feeding to satiation (Nie *et al.* 2017). This suggests that lethargy is not an inevitable result of feeding but an evolved response in some species. In behaviour regulation species both food intake and high energy reserve should lead to decreased physical activity, foraging behaviour and risk taking. Although anecdotally we generally experience lethargy after overfeeding or having substantial obesity, whether and to what extent humans develop lethargy after feeding is rarely quantitatively tested. Using meta-analysis of 11 overfeeding intervention studies in humans, we will demonstrate below that overfeeding leads to substantial decrease in physical activity and this decrease is largely responsible for the weight gain after overfeeding. We searched literature reporting human experiments which started with a baseline stable body weight in a study group and provided a constant increase in energy intake for a predecided period and then measured the change in weight. The change in food intake and body weight after the intervention should be related by the equation

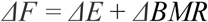

Where *ΔF* is the interventional change in energy intake, *ΔE* is the change in physical activity energy expenditure and *ΔBMR* the change in basic metabolic rate due to change in body weight. Basic metabolic rate (BMR) in humans is related to age, height, gender and body weight by the simplified Mifflin-St Jeor Equation (Mifflin *et al.* 1990) BMR (kcal/day) for males = 10 x Weight + 6.25 x Height – 5 x Age + 5 BMR (kcal/day) for females = 10 x Weight + 6.25 x Height – 5 x Age – 161 Since in the experiment the gender and height are constant and age can be considered practically constant over the experimental time, change in BMR should be directly proportional to change in weight, *ΔBMR = kΔW*. Since energy expenditure during a physical activity such as walking or running is also proportional to body weight, *ΔE* is also a function of *ΔW* such that *ΔE = pΔW*. Therefore,

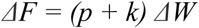

Since the experiments give data on *ΔF* and *ΔW*, and *k* can be estimated from the Mifflin-St Jeor Equation, *p*, the coefficient of change in physical activity can be calculated from the reported experimental data (Table 1). It can be seen that in all experiments ‘*p’* is negative and distributed over a narrow range with a mean of *-8.85* and standard deviation of *0.37*. This means that with a positive change in food intake the change in physical activity was always negative. Further if we calculate the resultant body weight if physical activity expenditure did not change with overeating, i.e. *p=0*, the increase in weight was substantially smaller. Therefore increased energy intake appears to have contributed only 11.5% and overfeeding induced lethargy 88.5 % to the increase in weight. It should be noted that the experimental intervention was only increasing the energy intake; no change in activity was intended, advised or suggested. However it appears to have followed overfeeding intervention naturally. Moreover the different independent experiments have a very consistent estimate of *p = −8.85*. This suggests that the decrease in physical activity on increasing food intake worked almost like a law. Thus the expectation of the hypothesis that lethargy should naturally follow food intake in humans is supported by very consistent evidence across independent studies. This meta-analysis also resolves the old debate of gluttony verses sloth (Prentice & Jebb 1995) by showing that it is gluttony induced sloth which is responsible for 88.5 % of weight gain across all overfeeding intervention studies.

**Table 1:** Meta-analysis of overfeeding intervention studies: All values have been converted to kcal for the sake of simplicity and following conversions have been used: One MegaJoule (MJ) = 239.006 kcal, One kilogram (kg) weight = 7716.17 kcal.
7. Adventure and aggression aversion in obese: Similar to physical activity, risk taking should be suppressed after food intake or on having high energy reserve according to our hypothesis. Conversely physical risk perception, physically aggressive or adventurous activities should facilitate CART-leptin interaction and thereby reduce adiposity. Some evidence in both the directions already exists. Obesity and anxiety have a positive association though the causality is unclear (Lykouras & Michopoulos 2011). Adventurous activities have been shown to boost weight loss (Jelalian *et al.* 2006). However, more rigorous tests of the prediction would be helpful. It is necessary to clarify here that in the context of evolution of mechanisms of foraging optimization, risk refers to the probability of being attacked, injured or struck by other wilderness associated dangers such as extreme temperatures etc. Health risks by alcohol, tobacco etc., are not included as risks in this context.
8. Physiological response to predator exposure: In a wide variety of species ranging from insects to mammals, presence of predator or stronger and aggressive conspecific competitor has been shown to reduce food intake, body weight or both (Blanchard *et al.* 1995; Carlsen *et al.* 1999; Higginson *et al.* 2012, 2016; Monarca *et al.* 2015; Ramler *et al.* 2014; Thaler *et al.* 2012; Tidhar *et al.* 2007). Although this has been generally interpreted as a “stress response” (Kestemont and Baras 2007) it is likely that it is a strategic response of the system evolved to optimize foraging gains against foraging risk (Tidhar *et al.* 2007).
9. Physiologically regulated versus behaviourally regulated species: In multispecies comparisons we expect that if kept in a safe captive environment with unrestricted food supply, the behavioural regulation species will become substantially more obese than the metabolic regulation species. Anecdotal and species specific accounts show that in certain species the differences in mean weights and adiposity as well as standard deviations between captive and wild animals are large (Schwitzer and Kaumanns 2001) and in certain species it is marginal (Chusyd et al 2018). A careful multi-species comparison can be used to test this prediction. At a proximate level we expect that metabolically regulated species would have a weak or no interaction of gut or adipose signals with risk and fear related signals. Leptin in these species should exert a food intake regulation effect with little influence of CART and other risk triggered peptides. Most studies on body weight regulation come from rodents followed by other animals including dog and primates all of which are behavioural regulation species. Our hypothesis predicts that leptin-CART and other interactions in elephant, horse and large ungulate species would be quantitatively or even qualitatively different than these species.
10. Further it is not necessary for the behavioural regulation hypothesis that food intake of obese individuals is greater than that of lean individuals. In human population weak positive correlation, lack of correlation and even negative correlations between obesity and energy intake are reported (Garaulet *et al.* 2000; Guillaume *et al.* 1998; Miller *et al.* 1990; Must & Tybor 2005; Wang *et al.* 2003) and all of them are compatible with the model.

| S. No. | PAPER NAME | YEAR | SUBJECTS | NUMBER OF SUBJECTS | PROTOCOL | kg/kcal | "p" |
| --- | --- | --- | --- | --- | --- | --- | --- |
| 1 | Diaz et al * | 1992 | Lean male | 7 | 7 months study divided in 5 periods - baseline (3 wk), overfeeding (6 wk), free diet, underfeeding and free diet. | 0.000119994 | -8.92 |
|  |  |  | Overweight males | 3 |  |  |  |
| 2 | Bouchard et al, Tremblay et al | 1990, 1992 | 12 pairs of identical twins | 24 | 2 periods - Baseline (14 days) and overfeeding (100days). For 84/100 days | 9.64286E-05 | -8.65 |
| 3 | Dallasso and James | 1984 | Normal males | 8 | 2 study periods of 7 day each - Baseline and overfeeding | 0.000106957 | -8.78 |
| 4 | Forbes et al | 1986 | Males | 2 | Maintenance period of 7 days followed by overfeeding period for max 3 weeks | 0.000148883 | -9.13 |
|  |  |  | Females | 13 |  |  |  |
| 5 | Glick et al | 1977 | Females normal | 4 | 2 periods of 5 days each - Baseline and overfeeding, followed by LCD for 4wks followed by 2 days of OF | 0.000155652 | -9.167 |
|  |  |  | Females obese | 4 |  |  |  |
| 6 | Jebb et al * | 1996 | Lean men | 6 | 7 days in metabolic suit followed by 12-d imposed mixed diets (3 overfeeding and 3 underfeeding) | 0.000155559 | -9.37 |
| 7 | Joosen et al | 2005 | Females | 14 | 2 periods - Baseline (7 days) and overfeeding (14 days). | 7.73826E-05 | -8.32 |
| 8 | Pasquet et al | 1992 | Males | 9 | 2 periods - Baseline (10 days) and overfeeding (61-65 days). | 7.44795E-05 | -8.26 |
| 9 | Ravussin et al | 1985 | Males | 5 | 2 periods - Baseline (13 days) and overfeeding (9 days) | 0.000185723 | -9.302 |
| 10 | Norgan and Durnin | 1980 | Males | 6 | 2 periods - Baseline (2 wk) and overfeeding (44 days) | 9.24835E-05 | -8.59 |
| 11 | Roberts et al | 1990 | Males | 7 | 3 periods - maintenance diet (10 days), overfeeding (21 days) and ad libitum intake (46 days) | 0.000118 | -8.9 |
| * only overfeeding period has been considered for this paper |  |  |  |  |  | Average | -8.85355 |
|  |  |  |  |  |  | Stdev | 0.37448 |

## Implications for obesity control

With the rapidly worsening obesity pandemic it is necessary and urgent to resolve between alternative hypotheses for the origin of obesity since the control measures can be substantially different for different hypotheses. If the behavioural regulation hypothesis is true, obesity control will depend not on diet composition and total amount of physical activity but on normalizing the CART-leptin and other necessary signal cross talks. Deficiencies of certain environment-behaviour interactions is responsible for the altered signalling. Since it is not possible to go back to the hunter gatherer life-style, the behavioural deficiencies can be compensated by behavioural supplementation. The easiest available behavioural supplementation is in the form of sports that mimic the deficient hunter gatherer behaviours. Most active sport involve hitting, kicking, chasing, attacking, defending and risk taking in some form or the other. These activities are expected to stimulate the signalling pathways activated by the natural hunter-gatherer behaviours. If these signalling mechanisms are normal it may not be necessary to cognitively control diet composition and calorie intake, the intrinsic mechanisms would be sufficient to maintain a homeostasis. Exercise is well recognized as a pro-health activity but the current popular perception of it is a means to burn calories. In our view, appropriate type of games and exercises supplement the missing behavioural triggers for CART and other signal molecules and thereby activate the intrinsic homeostatic mechanisms. It is possible that the behavioural component of the game contributes to health more than the calories burnt. If this is true, we expect that different types of exercises have different metabolic and endocrine effects although they might burn the same amount of calories. This indeed is demonstrated by many studies (Cuff *et al.* 2003; Eriksson *et al.* 1997, 1998; Hawley & Gibala 2009; Klimentidis *et al.* 2011; Nybo *et al.* 2010; Richards *et al.* 2010; Shen *et al.* 2015).

We show here that for species facing significant foraging risk, mechanisms for foraging optimization would evolve and the net food intake will be mainly under the control of the foraging optimization mechanisms. In such species decoupling of feeding and foraging risk can result into substantial increase in body weight. Humans belong to this category of species and therefore the evolved mechanisms of food intake regulation fail to work in the modern human society where feeding is decoupled from foraging. At a proximate level the interaction between obesity signals and risk induced signals decides food intake and thereby body weight. Reduction in risk induced peptide levels below a threshold can lead to a dramatic rise in mean body weight or adiposity of a population as well as high variance around the mean. Since the resultant adiposity in the absence of risk signals is unprecedented in human evolutionary history, the physiological mechanisms are not adapted to this level of adiposity. As a result adverse metabolic effects may precipitate. The model is likely to bring in a substantial change in the current thinking on which most preventive medicine for non-communicable diseases is based. It is necessary to test the predictions of the model and accordingly make fundamental changes to health policies which are likely to benefit generations to come.

## List of abbreviations

**List of abbreviations:**
|  |  |  |
| --- | --- | --- |
| 1 | CART | Cocaine Amphetamine Related Transcript |
| 2 | IUGR | Intra Uterine Growth Restriction |
| 3 | T2D | Type 2 Diabetes |
| 4 | MSH | Melanocytes Stimulating Hormone |
| 5 | POMC | Proopiomelanocortin |

## Declarations

- **Ethics approval and consent to participate**
- Not applicable since the study is only a theoretical model and analysis of published studies.
- **Consent for publication**
- All authors have their consent for publication
- **Availability of data and material**
- Not applicable since the model and results are completely described in the manuscript itself
- **Competing interests**
- The authors declare that they have no competing interests.

## Acknowledgements

- Not applicable

## Authors’ contributions

MW initiated the study and all authors contributed to the development of the concept and the model. MW and UB wrote the manuscript. All the authors read and approved the final manuscript.

